# Climate change and extreme weather events to intensify West Nile virus activity

**DOI:** 10.64898/2026.10.28.762036

**Authors:** Desiree Andersen Bliss, Gracie Fischer, Kacey Ernst, Nathan Schumaker, Michael A. Riehle, Elizabeth Hilborn, Stephanie DeFlorio-Barker, Leigh Combrink

## Abstract

Vector-borne zoonotic diseases pose a substantive risk to public health worldwide, yet predicting outbreaks at scales useful for public health response remains a critical challenge. Transmitted by *Culex* mosquitoes, West Nile virus is the leading cause of arboviral disease in the United States, progressing to neuroinvasive disease in approximately 1% of infections. Arizona’s record-breaking 2021 outbreak, with 1,715 reported human cases and 127 deaths exemplifies this growing burden. Using a decade of mosquito surveillance data coupled with time-lagged weather patterns in a random forest framework, we developed spatial models predicting *Culex* abundance and West Nile virus occurrence across Arizona at weekly temporal and fine spatial resolution. Our weighted models classified viral occurrence with 98.9% accuracy and explained 65.3% of weekly variation in county-level human cases, with human cases lagging modeled activity by one week. Neither mosquito abundance nor viral occurrence declined at the extreme temperatures characteristic of Arizona summers, suggesting that thermal refugia sustain transmission. Drought-monsoon cycling, rather than temperature alone, emerged as the primary driver of West Nile activity, and projections through 2100 indicate intensifying activity under all climate change scenarios.

## Introduction

Arboviruses, a subset of vector-borne pathogens transmitted by blood-feeding arthropods such as mosquitoes, ticks, and sandflies, account for over 17% of global infectious disease cases and more than 700,000 deaths annually. West Nile virus, first identified in Uganda in 1937, has emerged as the leading cause of arboviral disease in the United States [1]. Within five years of its 1999 introduction into New York City [2], WNV spread throughout the contiguous United States and has established itself as an endemic pathogen [3]. The virus circulates enzootically among birds, with *Culex* mosquitoes bridging transmission to dead-end hosts like humans and horses. While approximately 80% of human infections are asymptomatic, 1% progress to neuroinvasive disease [4], which carries a 10-30% mortality rate [5] and predominantly affects older adults and immunocompromised individuals [6,7]. Over 7 million human cases have been estimated in the U.S. between 1999 and 2016 [8], with outbreaks often following climatic anomalies such as heatwaves or prolonged droughts [9].

Mosquitoes are among the most efficient vectors of human pathogens, with life cycles tightly coupled to environmental conditions [10–12]. Temperature regulates mosquito development rates, survival and viral replication; precipitation shapes oviposition habitat availability; and humidity influences adult survival [13]. These relationships are non-linear: while higher mosquito densities generally correlate with increased pathogen spread, flooding or extreme heat can reduce *Culex* species survival [14]. Anthropogenic environmental changes such as urbanization further amplify pathogen circulation [15] by creating edge habitats favoring mosquito breeding.

Arizona has emerged as a critical U.S. hotspot for WNV. In its record-breaking 2021 outbreak there were 1,715 reported human cases and 127 deaths, representing about 59% of all U.S. cases that year [16,17]. The state’s arid climate, characterized by extreme heat and seasonal monsoons, creates ideal conditions for *Culex* mosquito proliferation: summer temperatures regularly exceeding 38°C (100°F) can speed mosquito development, increase biting rates, and shorten the extrinsic incubation period of the virus within mosquitoes [18,19]. Monsoon rains from July through September create ephemeral breeding sites while artificial water sources, such as irrigation canals and poorly maintained residential pools, sustain populations year-round [16,20]. Social, cultural, and economic disparities exacerbate Arizona’s WNV burden – low-income neighborhoods contain higher concentrations of neglected water sources [13,21], and outdoor workers face heightened vector-human contact during peak mosquito feeding hours [22,23].

Climate change is projected to intensify Arizona’s WNV burden through advancing temperature regimes and increased frequency of extreme weather events, particularly the drought-monsoon cycles that drive WNV activity [24,25]. Predictive models integrating climatic, ecological, and socioeconomic data are therefore urgently needed. Current modeling approaches are constrained by fundamental limitations: they focus solely on broad climatic patterns rather than fine-scale weather dynamics, lack operational forecasting capability or transferability due to reliance on site-specific data, or have insufficient accuracy for public health decision-making.

Here we present an integrated modeling framework that addresses these limitations. Our approach uses random forest algorithms to integrate a decade of mosquito abundance and WNV pool-positivity surveillance records with high-resolution time-lagged weather (representing biologically relevant processes) while also using a weighted model-averaging approach to reduce overfitting. We modeled spatial patterns of Arizona’s two primary WNV vectors – *Culex quinquefasciatus* and *Cx. tarsalis* – and WNV occurrence (pool positivity of mosquito traps). Our models achieved 98.9% classification accuracy for WNV occurrence in Arizona and explained 65.3% of weekly variation in human cases by county. We identified drought-monsoon cycling, specifically prolonged seasonal drought followed by heavy rainfall, as the primary driver of WNV activity and projected future outbreak patterns under multiple climate change scenarios. This work constitutes a transferrable and extensible, fine-scale vector-borne disease forecasting methodology designed for anticipating global zoonotic disease dynamics at weekly time scales.

## Results

### Predicting West Nile activity: from landscapes to human cases

Our WNV occurrence models, created using weighted random forest algorithms trained on 652,897 mosquito trap records collected between 2013 and 2023, achieved high classification performance (98.9%), maximizing both sensitivity (true positive rate; 98.95%) and specificity (true negative rate; 99.95%). West Nile occurrence probability was influenced primarily by the maximum temperature average recorded 0-5 weeks before sampling, followed by abundance predictions for both *Culex* species. When we aggregated predictions across all weeks to create a risk map index (**Figure 1C**), we identified concentrated risk at urban perimeters, particularly in Maricopa County, which consistently accounts for the majority of Arizona’s reported WNV cases and carries among the nation’s highest per-capita incidence [16,26,27].

**Figure 1.**
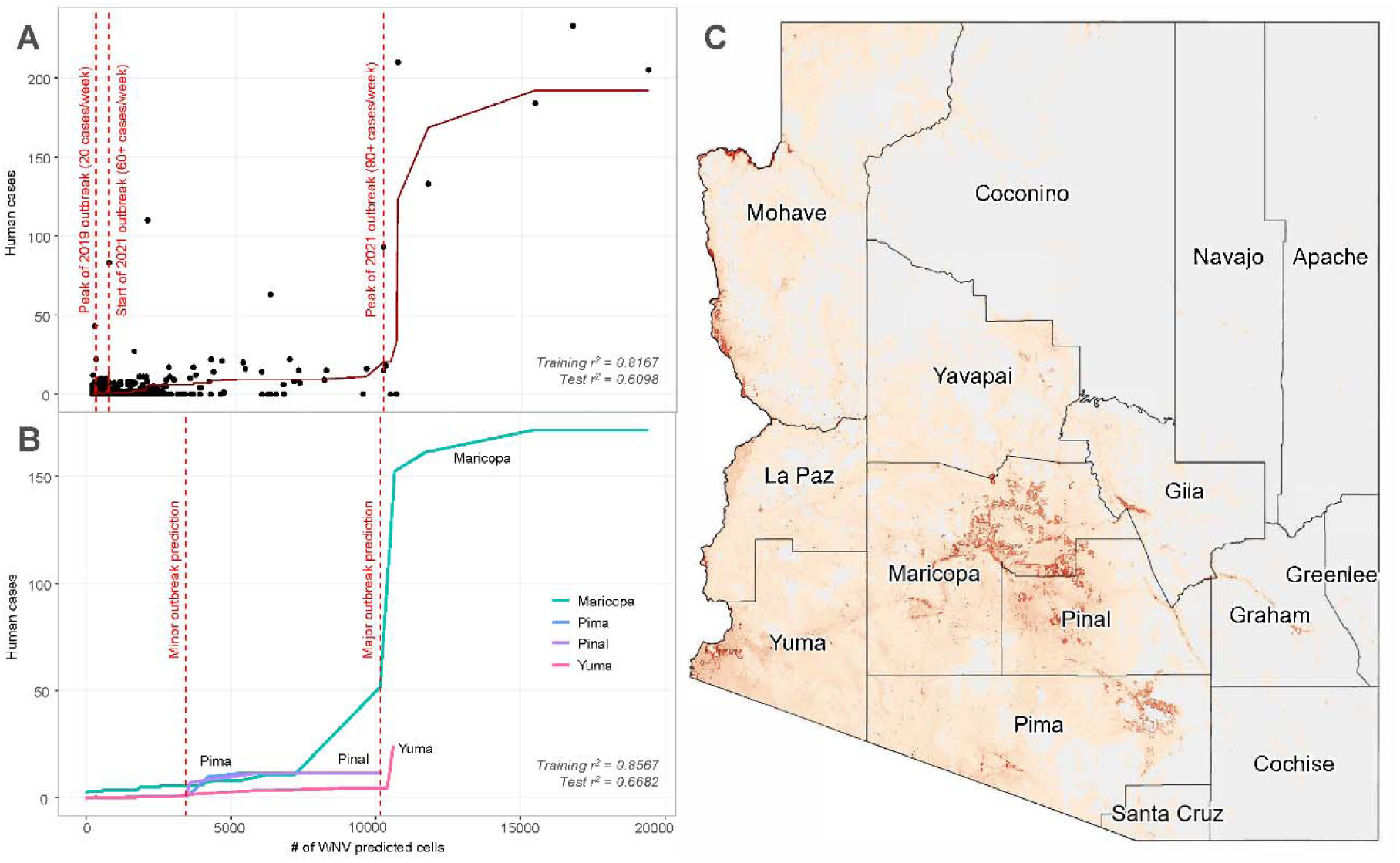
Human WNV cases and WNV risk map. Per-county numbers of human WNV cases predicted without (A) and with (B) county data used as a model covariate (vertical lines indicate significant outbreak events), and a map of WNV risk for Arizona across the study period (C).

The fit between modeled WNV activity and observed human cases provided real-world validation of our modeling approach. We explained 65.3% of the weekly variation in county-level human cases (2013-2023) using the area covered by WNV positive model cells (r^2^ = 0.6527) (**Figure 1A**). Incorporating county as a covariate increased explanatory power to 76.2% (r^2^ = 0.7617) and revealed two break points for minor and major outbreaks (**Figure 1B**). Critically, modeled WNV activity was most strongly associated with human cases one week later, representing a comparable time frame to the time between mosquito exposure and symptom development (2-6 days, up to 14 days)[28].

### Mosquitoes and West Nile virus are not limited by extreme high ambient temperatures

Temperature emerged as a primary driver for WNV occurrence and both *Culex* species’ abundance with species-specific temporal patterns reflecting differing biological adaptations. For *Cx. quinquefasciatus* (r^2^ = 0.3262), mean temperature average 0-3 weeks before sampling showed greatest influence, indicating rapid demographic responses to immediate thermal conditions. In contrast, *Cx. tarsalis* (r^2^ = 0.4445) responded most strongly to minimum temperature 5-13 weeks prior, suggesting population dynamics were influenced by early-season survival and development. Both species showed increasing abundance with rising temperatures, with no evidence of thermal limitation even at extreme high temperatures.

West Nile occurrence probability also grew with increasing maximum temperature of 0-5 weeks before trapping. Concerningly, while critical thermal maxima are known to inhibit mosquito survival [29,30] and within-mosquito WNV replication (CTmax up to 38.6°C in *Cx. tarsalis*) [31], the empirical data did not show decreases in modeled mosquito abundance or WNV probability at extreme high temperatures (**Figure 2A-C**). This could be due to infected mosquitoes utilizing natural and man-made microhabitats which allow both mosquito and virus to escape the extreme high temperatures of Arizona summers. Alternatively, these patterns may indicate adaptation has occurred over time.

**Figure 2.**
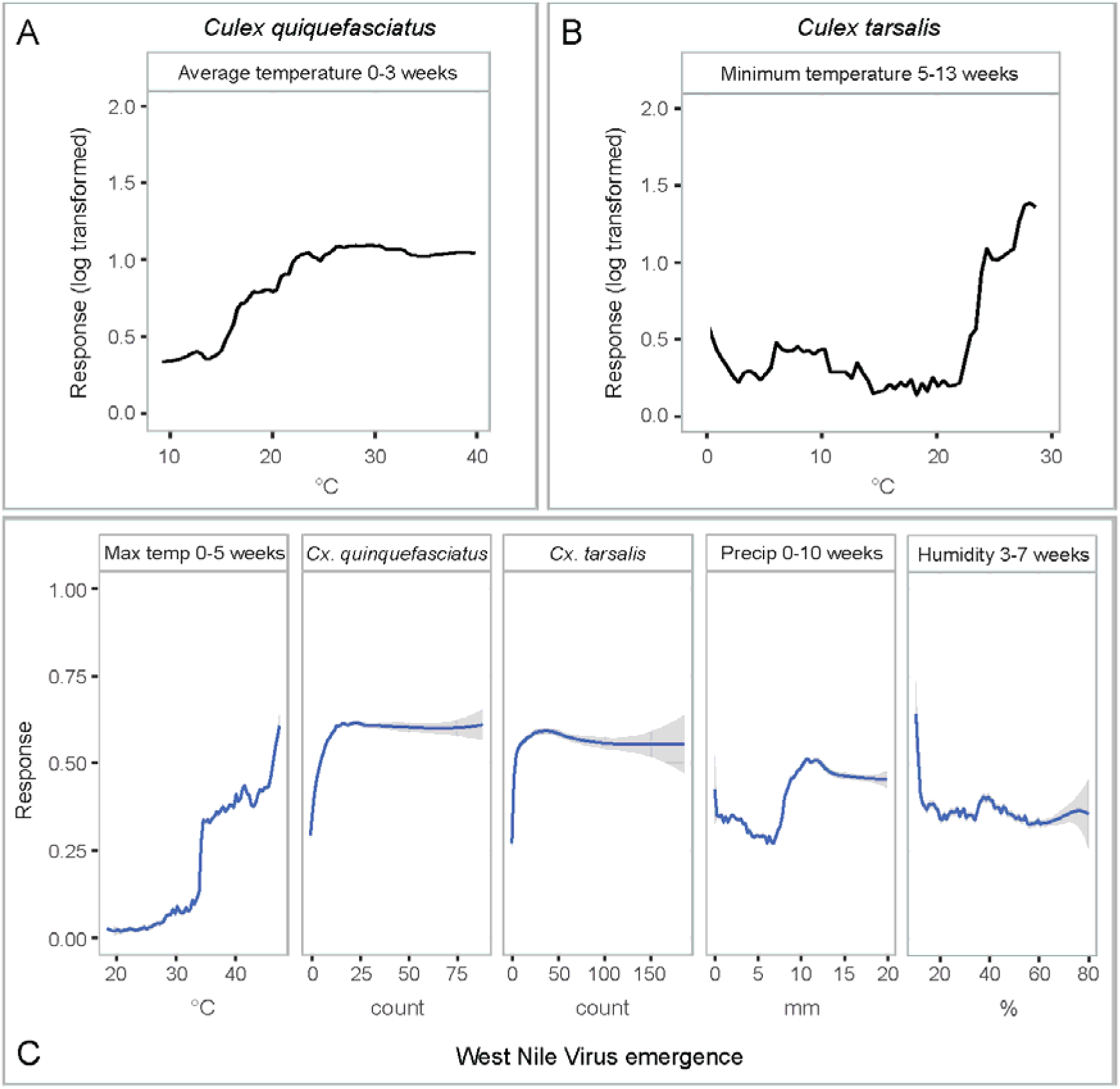
Variable response curves and standard error. Highest importance variable responses for *Culex quinquefasciatus* (A), *Cx. tarsalis* (B), and West Nile virus activity (C). Response curves for WNV are arranged from highest importance (left) to lowest importance (right). Response curves for all variables for both *Culex* species models available in Figures S2 and S3, and all West Nile virus variable responses in Figure S5.

### Cycles of drought and monsoon exacerbate WNV on the landscape

Our WNV occurrence models revealed the critical importance of drought-monsoon cycling in driving viral transmission, with the selected variables reflecting seasonal patterns endemic to the state of Arizona (**Figure 2C**). Maximum temperature 0-5 weeks before trapping events, the highest importance variable in our occurrence model, captures the extreme heat associated with pre-monsoon drought conditions. Similar to our mosquito species’ abundance, WNV activity increased with temperature and did not appear to drop off at the most extreme high temperatures of 47.4°C (maximum temperature averaged for 0-5 weeks within trapping), or about 117°F, corresponding with heat waves and potential drought. Similarly, we saw the highest peaks in WNV activity happening below 15% (extremely dry) and around 35% (dry) humidity during the 3-7 weeks before sampling.

Human WNV cases in Arizona peak around calendar week 34 (week 34.1 ± 4.5; data from ADHS), or the 3rd week of August, corresponding to the 10th week of monsoon season, which starts on June 15. In our models, viral activity peaked at 10 mm of precipitation 0-10 weeks before mosquito collection and tapered off slightly at higher precipitation levels. This indicates that wetter than average monsoons do not greatly increase WNV, but rather a typical monsoon acts as a catalyst for activity. Our predicted peak in WNV activity matches with average weekly precipitation during monsoon season in the Phoenix metropolitan area (∼10.8 mm), with Sedona (∼16.7 mm), Flagstaff (∼29.0 mm), and Tucson (∼29.7 mm) falling within similarly high risk for their respective weekly monsoon precipitation levels. Yuma, the state’s third largest metro area, has an average weekly monsoon precipitation of ∼3.2 mm, putting it below the peak observed in our models. This could be why Yuma County had such low numbers of human WNV cases – only 8 cases have been reported between 2013-2023, out of 2,526 cases statewide.

### Climate change induced extreme weather variability may intensify WNV activity with regional variation

Our climate change projections suggest complex WNV activity patterns for the future of Arizona. Using two climate models spanning a range of equilibrium climate sensitivities (INM-CM5-0 [32] with 1.9°C ECS; UKESM1-0-LL [33] with 5.4°C ECS) and four shared socioeconomic pathways (SSP126 the “sustainability” scenario; SSP245 the “middle of the road” or “regional rivalry” scenario; SSP370 the “inequality” scenario; and SSP585 the “fossil-fueled development” scenario [34]), we projected activity patterns through 2100. By analyzing trends across the state, we determined that significant increases in number of weeks of viral activity would occur across 0.6% to 39.4% of the state depending on climate model and scenario, while decreases would occur across 0.1% to 0.3% of the state (**Figure 3**). Under the assumptions of the UKESM model, the most extreme climate change, by 2100 parts of the state could see 10 extra weeks of WNV activity. The greatest increases will likely occur at higher altitudes, where lower temperatures have historically limited viral transmission. Interestingly, some areas around the Phoenix metropolitan area and Yuma are expected to see a decrease in the number of weeks of activity possibly due to changes in precipitation patterns.

**Figure 3.**
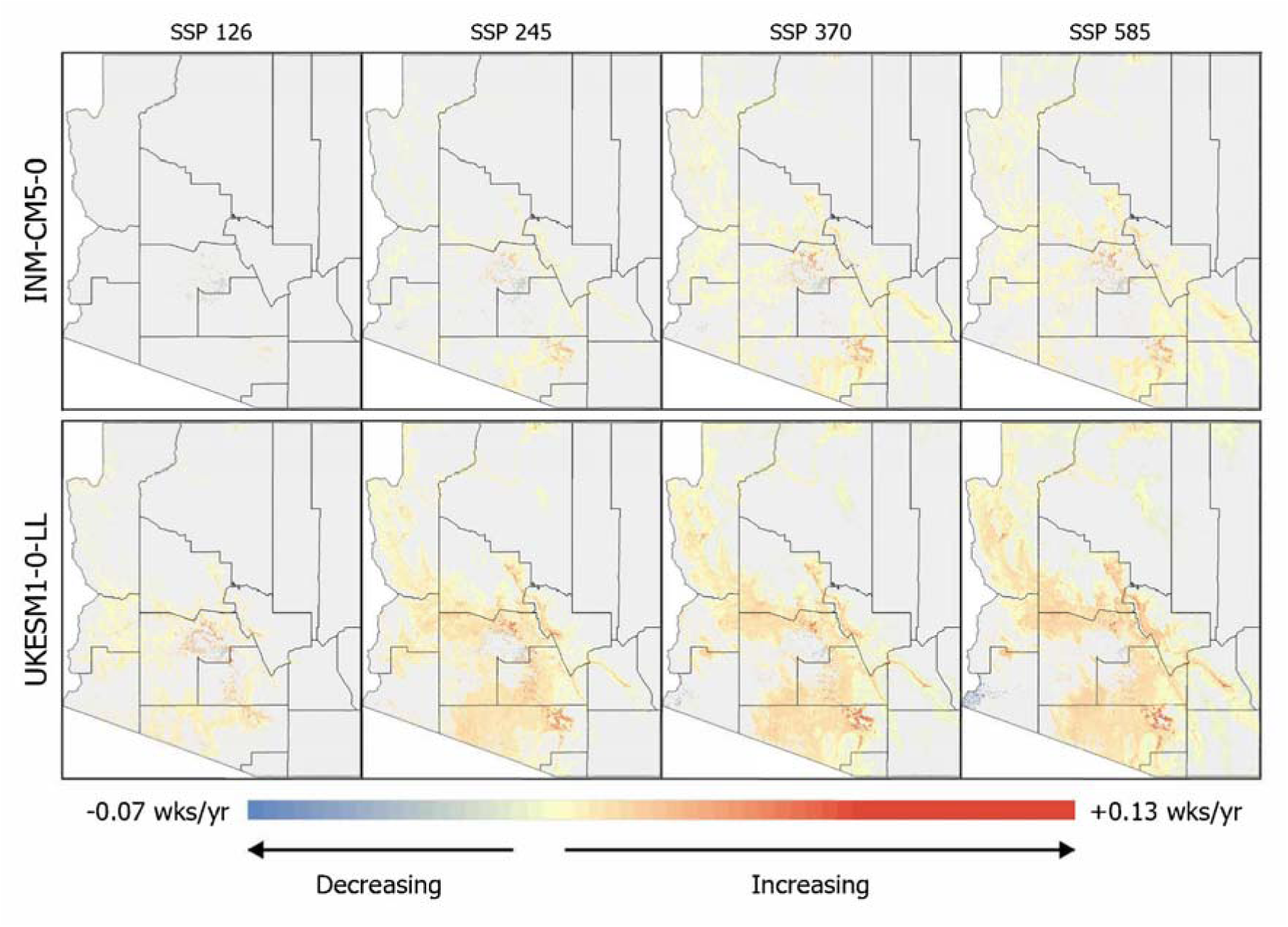
Climate change trends. Maps of significant trends in the number of weeks of West Nile virus activity in Arizona under four climate change scenarios from two climate models (low vs. high sensitivity). SSP126 represents the “sustainability” scenario, SSP245 the “middle of the road” or “regional rivalry” scenario, SSP370 the “inequality” scenario, and SSP585 the “fossil-fueled development” scenario.

## Discussion

The lack of a population size constraint on *Culex* species imposed by high temperatures has profound implications for climate change impacts. Rather than being limited by extreme temperatures, mosquito and viral populations may continue expanding as thermal refugia remain available. The increasing prevalence of urban heat islands and irrigation likely expand these refugia, further supporting mosquito and viral persistence on the landscape.

The patterns we observe in our models bolster the hypothesis that drought and monsoon cycles drive WNV activity [24]. Drought creates limited water sources, which in turn concentrates both avian hosts and mosquito vectors in small areas of close proximity [25]. Stress from heat and limited water then amplifies WNV transmission and replication. When the subsequent monsoon rains fall, mosquito populations can rapidly increase, facilitating pathogen transmission to humans. This drought-monsoon cycling explains Arizona’s distinctive epidemiology and provides a framework for understanding disease transmission in arid regions around the world.

Climate change will further exacerbate not only total activity weeks, but also vector abundance and the intensity of WNV transmission. Rising temperatures extend mosquito breeding seasons, and irregular rainfall patterns create ephemeral pools ideal for larval development [35]. For instance, increases in average temperatures can reduce *Culex* larval development time, thereby accelerating viral transmission cycles [36]. In our projections, the greatest increases will likely occur at higher altitudes, where lower temperatures have historically limited viral transmission. Similar expansions into altitudinal gradients have already been seen with malaria [37] and dengue [38].

The projected decreases in activity around the Phoenix metropolitan area and Yuma are likely linked to decreasing rainfall with anticipated climate change. Yuma is already well below the precipitation threshold of peak WNV activity (3.2 mm in Yuma vs. 10 mm optimum), and parts of the Phoenix metropolitan area will likely fall below this threshold under some climate change scenarios. Importantly, our projections incorporate only direct climatic effects on WNV activity patterns. Confounding factors such as urbanization, water management, mosquito adaptation, and viral evolution may further affect future transmission patterns. For example, extensive irrigation systems across Phoenix and Yuma will continue to provide unnatural water sources outside of the monsoon and decreasing precipitation.

The methodological advances described here set a baseline for a novel transferable high-resolution (spatial and temporal) disease forecasting model framework. The integration of time-lagged weather variables in a random forest framework captures biological processes that contribute to vector abundance and viral activity. Using weighted model averaging, we reduce overfitting while maintaining sensitivity to complex environmental interactions. Critically, we demonstrate real-world relevance by correlating our models with human WNV case data at a one-week time lag, which makes our predicted activity patterns epidemiologically meaningful.

The transferability of our framework extends beyond WNV and Arizona; all predictor variables are publicly available and can be incorporated into models of other vector-borne disease systems and geographic ranges. Additionally, the fine temporal resolution of our models is what makes this framework relevant to public health decision-making, as managers can identify in real-time where disease outbreaks may occur, facilitating enhanced surveillance and early case detection. For operational implementation, our framework requires surveillance networks of vectors and pathogens, which are already implemented in many parts of the United States and around the world. Once the modeling framework is calibrated for the specific system and geographic area, weekly predictions can be automated to provide early warning systems for public health agencies, healthcare providers, and even the general public. These advanced forecasting models could then inform mitigation strategies that target larvicide application, public education campaigns, and healthcare system preparation.

The potential public health benefits of spatial disease forecasting are already substantial, and growing. Current vector and disease surveillance are primarily constrained to reactive mitigation from case reporting after disease activity, limiting timely intervention. The development of our modeling framework was designed with transferability in mind. It is our hope that our models will be adapted to address other endemic and emerging threats, including St. Louis Encephalitis Virus (SLEV), and *Aedes aegypti*-borne pathogens like Zika and dengue in the US, with potential application for diseases like malaria in tropical regions. This work is timely, as our framework shifts this paradigm to predictive forecasting and enables preemptive strategies rather than reactive ones, thereby allowing for potentially significant reductions in morbidity and mortality while decreasing the burden on healthcare systems during outbreaks.

## Material and methods

### Random Forest Algorithms for Spatial Modeling

We selected Random Forest (RF; ‘randomForest’ package in R [39]) as our modeling algorithm due to its accuracy and ability to detect complex relationships between predictor and response variables. It is also a form of model averaging, which uses all decision trees in the forest to form a prediction based on agreement of all trees. Unlike traditional statistical approaches, RF handles high-dimensional datasets with collinear variables, missing data, and nonlinear relationships [40]. This flexibility makes RF ideal for modeling complex systems like mosquito-borne disease transmission.

RF’s ability to rank variable importance offers unique insights into disease drivers. In dengue forecasting, RF identified rainfall, temperature, and urban mobility as critical predictors at long- and short-term forecasting horizons [41]. By isolating influential variables, RF helps prioritize public health interventions in resource-limited settings. RF’s capacity to synthesize satellite imagery, groundwater use, and socioeconomic indicators provides a scalable framework for generating actionable risk maps. Additionally, RF’s robustness to noise and outliers enhances its utility in real-world applications [42].

Random forests are uniquely suited to modeling mosquito-borne diseases due to their ability to integrate diverse data types. For example, Chen et al. [43] combined socioeconomic variables (e.g., income, education) with landscape metrics (e.g., vegetation cover, water body density) in a k-nearest neighbor (kNN) model to predict mosquito abundance in Charlotte, NC, achieving an accuracy of 95%-99%. This approach revealed that heterogeneous landscapes with mixed residential and green spaces harbored higher mosquito densities. These results are intended to guide future urban planning in an effort to reduce mosquito habitat availability. In South Korea, an RF model using land-use and meteorological data achieved the highest predictive accuracy across multiple urban clusters, which highlights its robustness in handling complex ecological data [44]. In Colombia, RF models forecasting dengue outbreaks at sub-national scales incorporated satellite-derived land surface temperatures and education coverage, outperforming artificial neural networks and traditional statistical approaches and demonstrating strong predictive performance across short- and long-term time scales [41]. These studies demonstrate RF’s ability to capture complex interactions between environmental and anthropogenic factors.

### Mosquito Surveillance Dataset

In Arizona, the Department of Health Services (ADHS) aggregates county-level surveillance data of mosquito populations year-round with the majority of active surveillance occurring from May-September. County-level trapping varies significantly across Arizona; the highest trapping is conducted in Maricopa County where the Maricopa County Vector Control Program places 800 traps weekly throughout the year. Other counties have modest levels of trapping, with some counties placing no or minimal traps exclusively during the mosquito season. Some trapping is conducted as strictly surveillance, while other trapping is conducted in response to mosquito exceedances or West Nile virus positivity. Despite differences in geographic and temporal coverage, the trapping methods are similar. Briefly, CO_2_ mosquito traps, baited with dry ice, are placed at designated locations across a grid with additional trap locations selected based on previous indications of high mosquito activity and varying by county [45]. Traps are set in the evenings and left out overnight and collected the following morning. Mosquitos are morphologically identified to the species level, with target *Culex* species then being transported to either local county health departments or the Arizona State Public Health Laboratory for testing via Rapid Analyte Measured Platform (RAMP). Pools of target *Culex* species that test positive for West Nile virus (WNV) via RAMP are sent to Maricopa County for confirmation of WNV presence using polymerase chain reaction (PCR) tests.

We obtained statewide mosquito surveillance data spanning from January 3, 2013, to September 29, 2023, from the Arizona Department of Health Services. These data were compiled and simplified to include the date collected, latitude, longitude, species, number of mosquitoes collected, sex, arbovirus tested, and arbovirus case status. Year and week were extracted from the collection date and concatenated to obtain a temporal identifier. The compiled dataset included 652,897 data points collected across 543 weeks from 8,310 unique trap locations throughout all 15 Arizona counties. A total of 21 mosquito species were recorded. These data also included 233,807 absence points, where no mosquitoes were collected. This dataset was then divided into subsets for the two main mosquito vectors in Arizona, *Culex quinquefasciatus* (103,434 collections with 516,726 collected) and *Culex tarsalis* (37,180 collections with 377,844 collected), as well as one subset for WNV occurrence (129,389 traps, 2,846 positive traps, 153,224 pools tested, 3,019 positive pools). For mosquito abundance, we aggregated female abundance only. *Culex* abundance was log-transformed.

### Environmental Data

Weather data were obtained from 47 Arizona weather stations from the Automated Surface Observing System (ASOS) network available through the Iowa State University’s Iowa Environmental Mesonet (IEM)[46]. For the period January 3, 2013, to September 29, 2023, we downloaded hourly data on temperature, precipitation, and humidity. These data were aggregated for each station, by year and week. Minimum, mean, and maximum temperature in degrees Celsius were computed per week, and accumulated precipitation was calculated in mm. Humidity was averaged for each week. Weekly surface weather map layers were derived using the Empirical Bayesian Kriging (EBK) Regression Prediction tool (ArcGIS Pro, ESRI, Redlands, CA), with a digital elevation model (Airbus, USGS, NGA, NASA, NOAA, CGIAR, GEBCO, NLS, OS, NMA, Geodatastyrelsen, GSA, GSI and the GIS User Community) serving as the explanatory variable. Environmental layers used as predictors for training the models had a resolution of 2.7 x 10^-4^ decimal degrees (dd), i.e. roughly 27 meters. Custom R functions were used to link the mosquito surveillance data points to the weather layers using time lags that ranged from 0 to 20 weeks. These time lagged weather data were then smoothed by averaging for intervals of 2-21 weeks, to represent biologically and phenologically important weather patterns such as temperature, which has been shown to correlate with mosquito species abundance[47]. In total, we had 1,155 time-lagged and averaged weather variables for minimum, mean, and maximum temperature (tmin, tavg, tmax, respectively), total precipitation (prec), and average humidity (humd). In addition, we developed averaged indices for both the U.S. Geological Survey Landsat-8 NDVI (normalized difference vegetation index) and MNDWI (modified normalized difference water index [48]) for the study period, for all imagery with <10% cloud cover. Finally, we used World Land Cover 30m BaseVue 2013 (MDAUS, underlying data updated in 2017) to represent landcover classes. This older source data was selected due to its ability to identify landcover types at fine scales, while still remaining accurate when compared to more recent satellite imagery.

### Modeling

For *Cx. quinquefasciatus* and *Cx. tarsalis*, we used randomForest for regression of log-transformed abundance. For WNV, we used randomForest for classification of positive versus negative pools (Figure S1). Also for WNV, we divided the data into positives and negatives, then randomly divided negatives into 44 subsets, maintaining an equivalent number of points to the positive set (2,845 positives in total). Each of these 44 subsets were then trained along with the positive set to create 44 separate random forest models. All models were stratified by county (randomForest function setting ‘strata’) to reduce the weight of county sampling effort on the models.

Variable selection involved creating models trained with subsets of the weather predictor variables and maintaining those variables with the highest importance, represented by the increase in node purity (Tables S1–S3). This selection process was repeated until a manageable number of variables remained; then, highly correlated variables with low importance were removed until none remained (Figures S2 and S3). Variables with the lowest importance were removed until their absence had a significant negative impact on the r-square of the final decision tree of the model. In the case of WNV, variables with the lowest importance were removed until the combined class error was significantly increased. For WNV, we compiled the selected variables from all 44 models and maintained variables that represented the majority of all models, removing highly correlated variables (Figure S4). This allowed us to maintain consistency among all models. Both *Culex* species models were extrapolated to the WNV dataset and included as predictor variables for WNV. To create surface rasters of *Culex* abundance and WNV presence, we projected our final models to all weeks between 2013-2023 and to each climate model, SSP, and year (16 projections). Layers used for projecting models had a resolution of 0.02 dd (∼2 km), a scale we used to reduce processing time and power.

### Future Projections

To determine the potential impacts of shifting weather patterns on mosquitoes and WNV, we extrapolated our weekly weather data using two climate models and four shared socio-economic pathways (SSPs). The INM-CM5-0 [32] climate model has a low sensitivity of 1.9 °C equilibrium climate sensitivity (ECS), while the UKESM1-0-LL [33] climate model has a high sensitivity of 5.4°C ECS. By using projections from both models, we were able to evaluate a range of possible outcomes capturing different levels of possible radiative forcing. We also used four SSPs within each climate model to represent scenarios associated with low (SSP126, SSP245) to high (SSP370, SSP585) forcing [34]. To predict weekly weather patterns for time periods between 2021-2040, 2041-2060, 2061-2080, and 2081-2100, we first extracted the monthly difference between the current and future predicted temperature and precipitation from Worldclim.org. These differences were then applied to weekly weather data for the relevant month.

### Human cases

Human WNV case data for the study period (2013-2023) was obtained from the Arizona Department of Health Services and number of cases were broken down by year, week, and county. No identifying information was obtained or used. Random forest was again used to model the number of human cases per week per county by the number of model predicted WNV positive cells within each county as calculated by the Zonal Statistics as Table tool in ArcGIS Pro. The number of cells was lagged to 0-6 weeks prior to the reported week of human cases, and each lagged week was used to create separate models. We selected the model with the highest r^2^ to represent an optimal lag to predict potential human outbreaks.

### Model Fit

Random forest models for the two *Culex* species were evaluated on average r^2^ of the forest and on r^2^ of a linear model of predicted versus actual value when the models were predicted to the full datasets. For WNV, we evaluated averaged sensitivity (true positive rate) plus specificity (true negative rate) across all 44 models. We also extrapolated all 44 models to the full dataset and created a sum of model agreement represented by the total number of predicted positives ranging 0-44. Finally, we thresholded these summed predictions at each value between 0 and 44 and compared them to actual values to determine the maximum sensitivity plus specificity (maxss) threshold and value.

### Curve Fitting Trends

To quantify trends in WNV activity, we calculated, for each year, the earliest and latest week of activity and the total number of active weeks. To relate observed dynamics to projected climate conditions, we assigned proxy years to climate projections by aligning decadal-scale weather patterns, for example, mapping observations from 2013-2023 onto the 2028-2038 period within the 2021-2040 projection window. Raster layers representing weekly activity metrics were converted to point-based datasets to enable location-specific analyses. At each point, we fit a linear regression of activity metrics against time and considered trends statistically significant at p < 0.05. For locations with significant models, slope coefficients were used to quantify the direction and magnitude of temporal change.

## Acknowledgements

We thank Jacob Hojnacki and Sierra A. Jaramillo from Coconino County Health and Human Services (Arizona, USA) and James Markweise of the US Environmental Protection Agency for their contributions during early stages of this work. We additionally thank Irene Ruberto and Guillermo Adame of the Arizona Department of Health Services for their cooperation and collaboration in sharing data.

The authors used a generative AI tool in the preparation of this manuscript. Model: Claude Sonnet 5.5 (Anthropic). Tasks delegated: proofreading and editing of author-written text. All AI-assisted outputs were reviewed, verified, and edited by the authors, who are solely accountable for the accuracy and integrity of the work. The AI tool is not listed as an author.

## Author contributions

Conceptualization: DAB, LC, KE, EH, NS, MR, SD-B; Methodology: DAB; Investigation: DAB; Visualization: DAB; Funding acquisition: LC, KE, NS; Supervision: LC; Writing – original draft: DAB, GAF, LC; Writing – review & editing: DAB, LC, KE, EH, NS, MR, SD-B

## Data, code, and materials availability

All mosquito surveillance data and climate projection data used in this analysis are available from the Arizona Department of Health Services and the sources described in the Methods. Code for the random forest modeling framework is publicly available at https://github.com/desireeka93/WNV-Activity [49].

