## Supplementary Information for "Climate change and extreme weather events to intensify West Nile virus activity"

Figure S1.

Methodology flowchart for mosquito and West Nile virus modeling in Arizona.

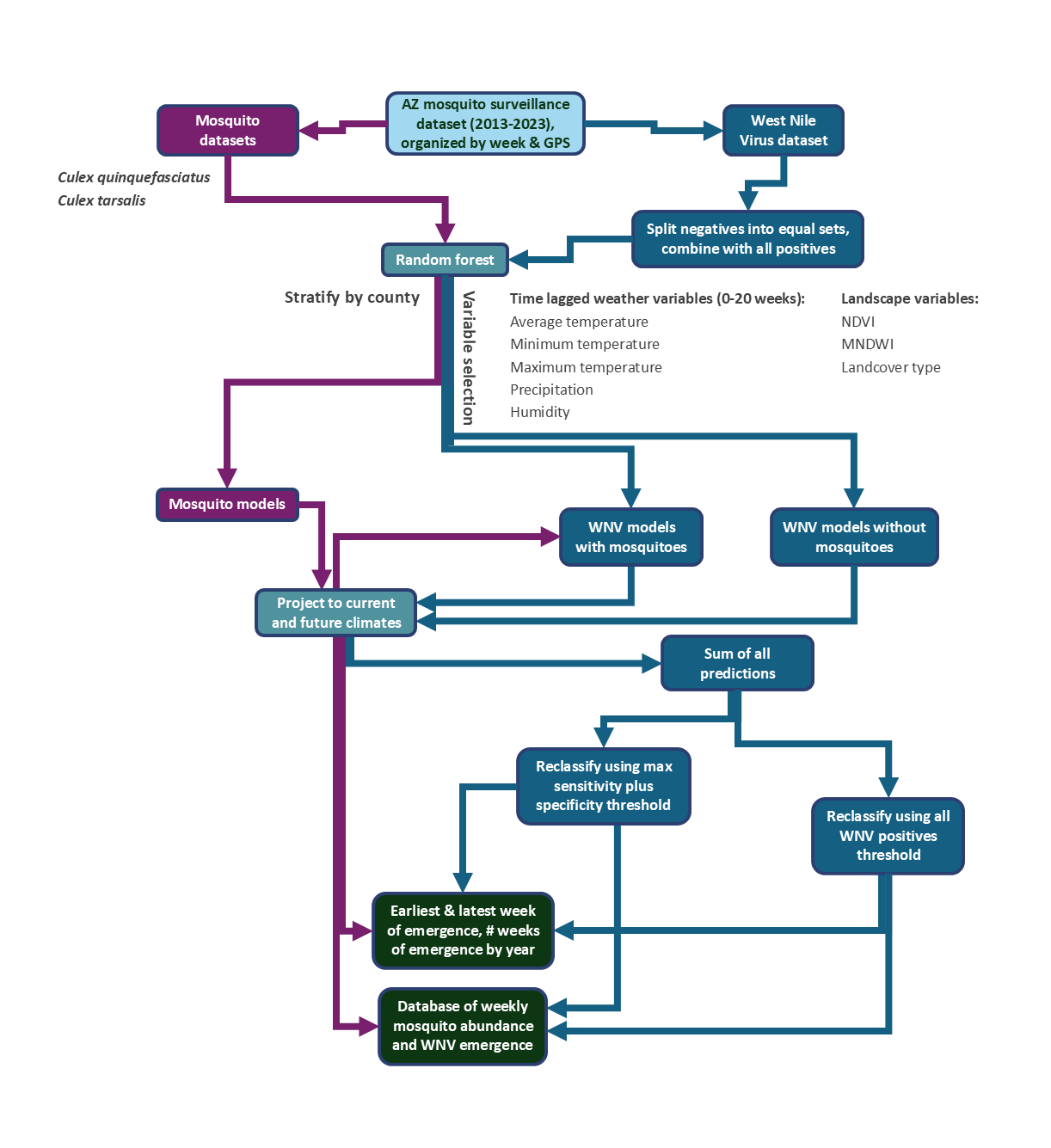

Figure S2.

Selected variables for *Culex quinquefasciatus* model.

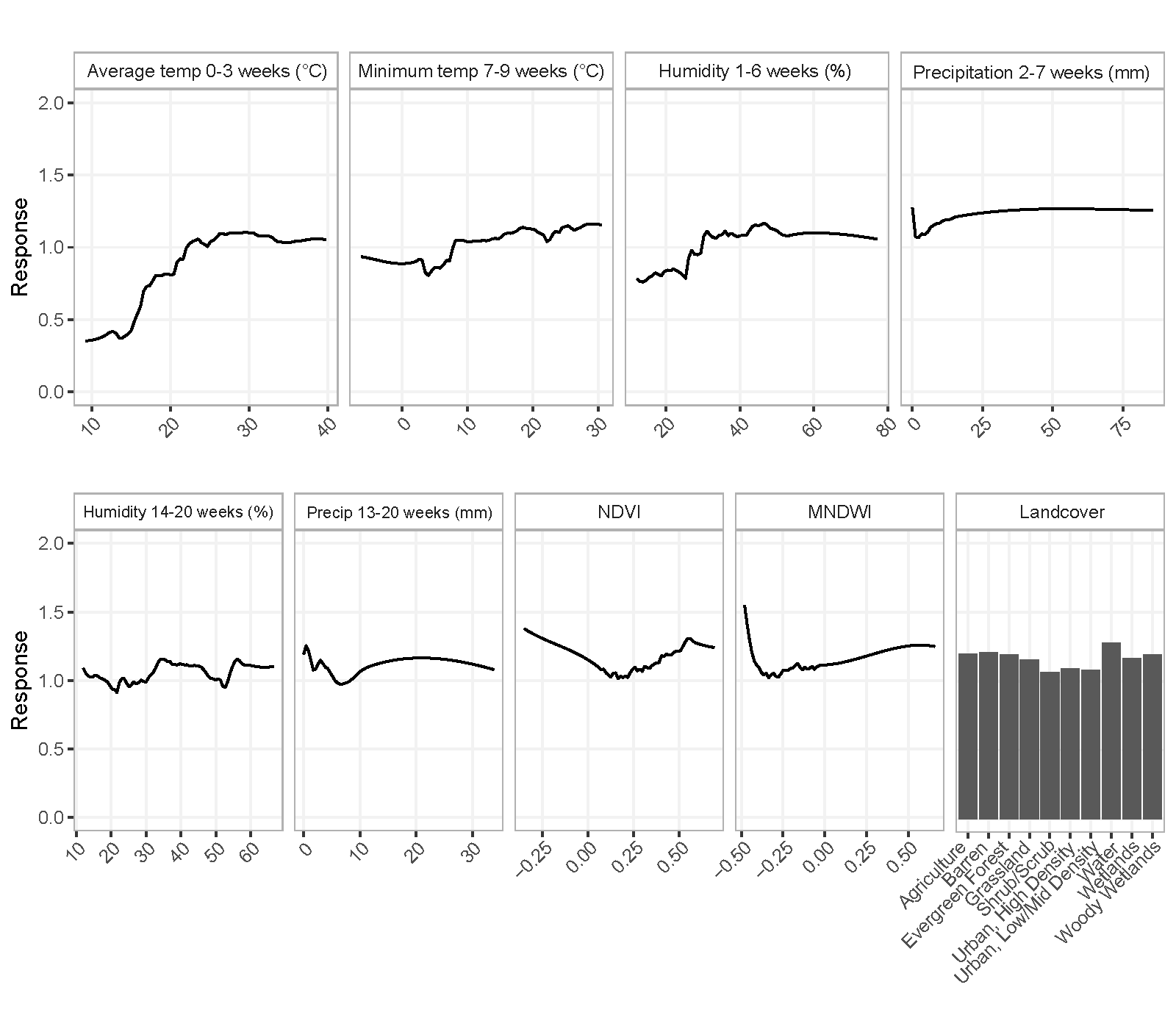

Figure S3.

Selected variables for *Culex tarsalis* model.

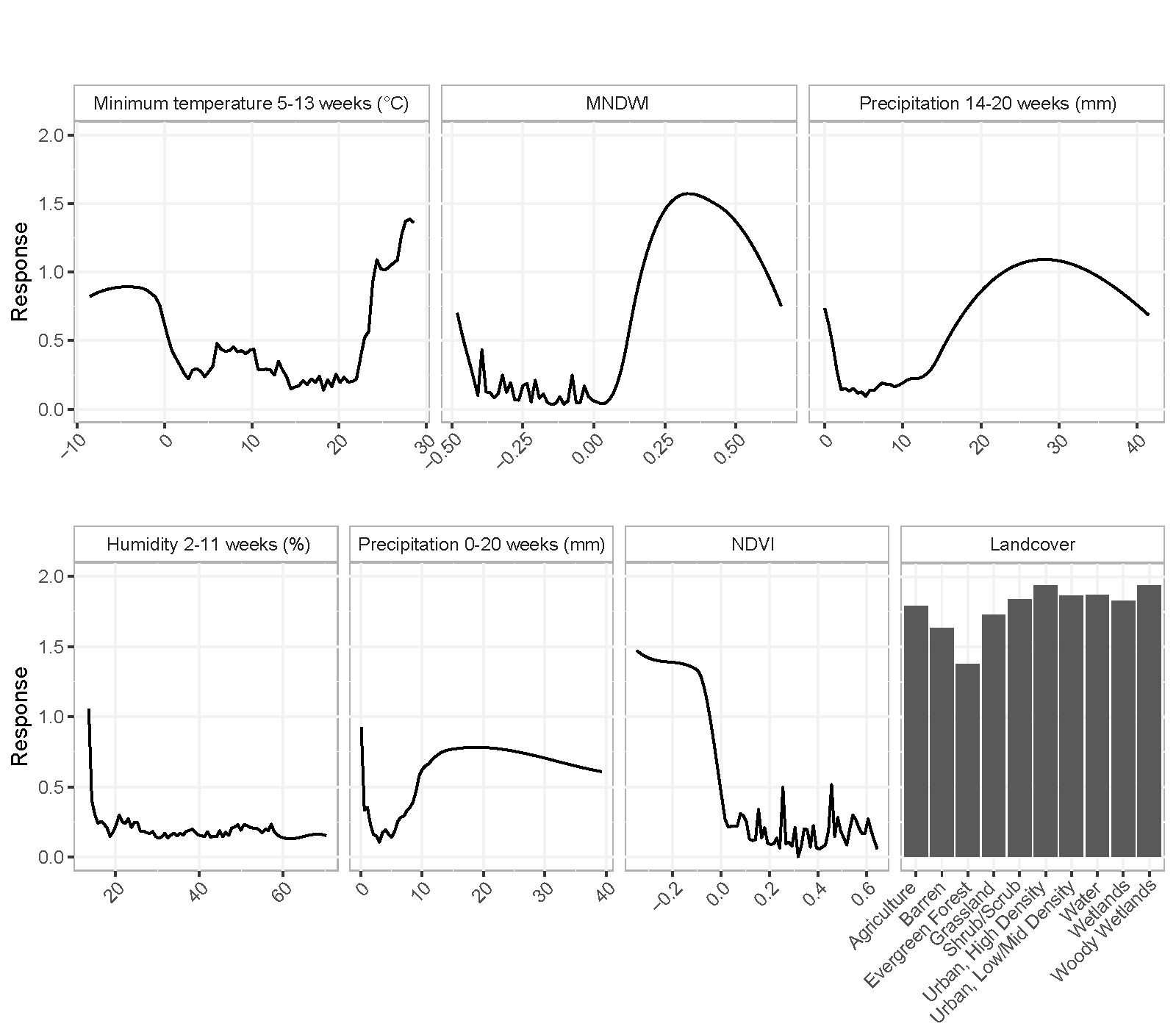

Figure S4.

Selected variable importance for West Nile virus model

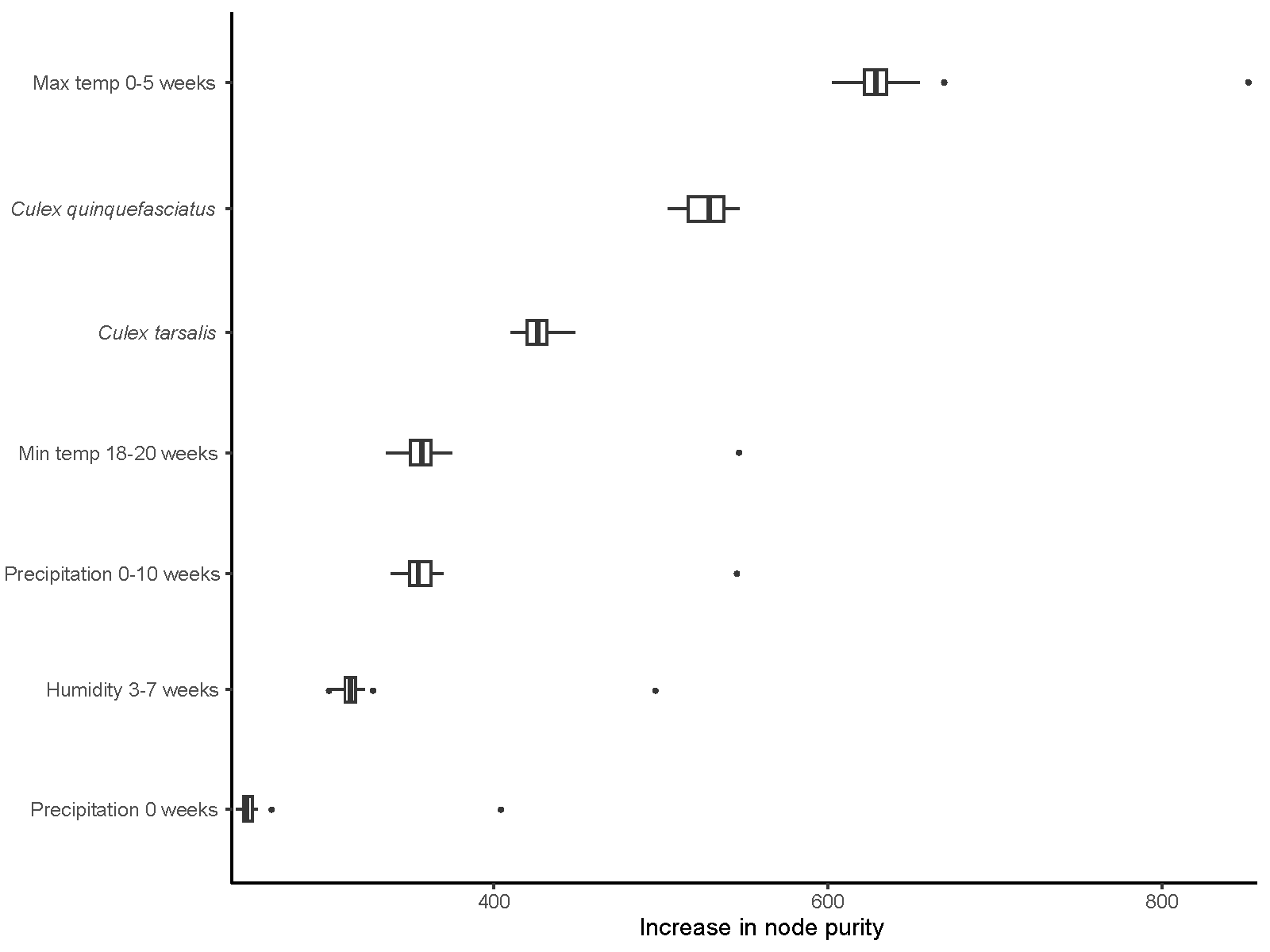

Figure S5.

All variable responses for West Nile virus model

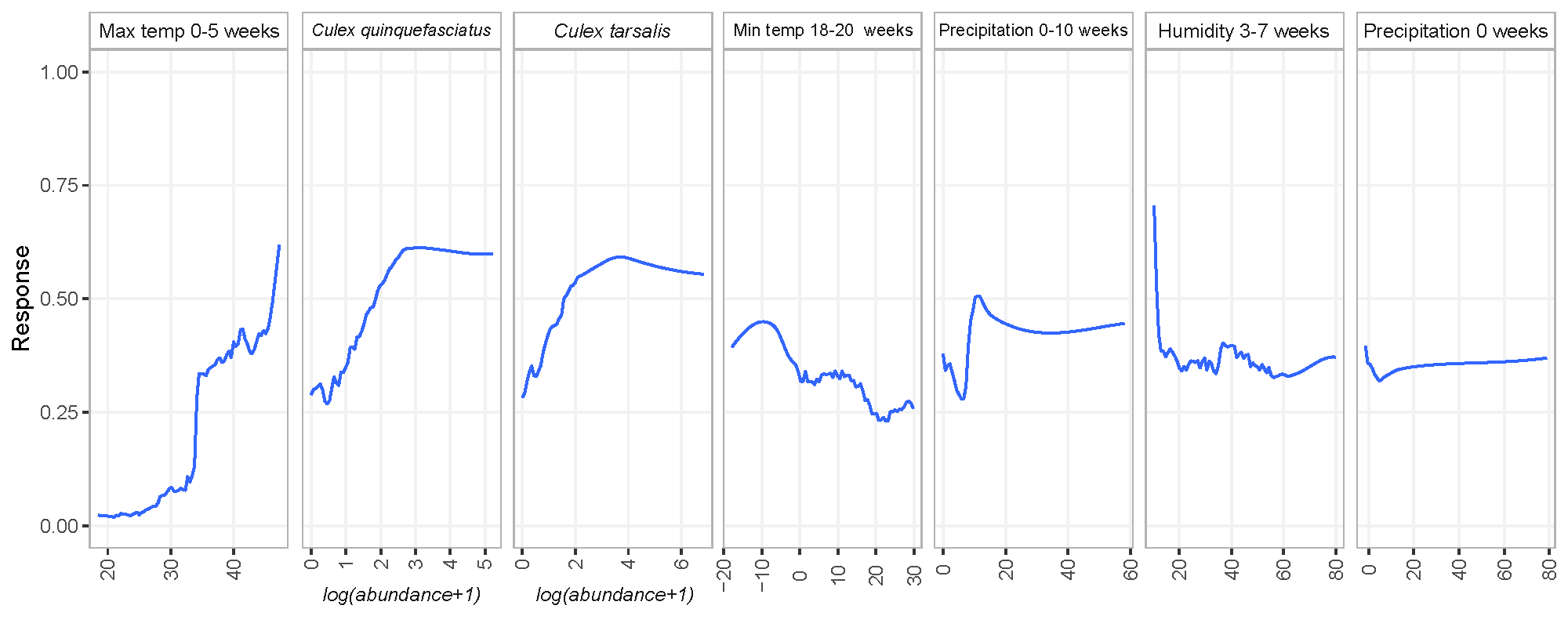

Table S1. (separate file)

Variable importance used for selection of variables in *Culex quinquefasicatus* model. Available at: <https://github.com/desireeka93/WNV-Activity>.

Table S2. (separate file)

Variable importance used for selection of variables in *Culex tarsalis* model. Available at: <https://github.com/desireeka93/WNV-Activity>.

**Table S3. (separate file)**

Variable importance used for selection of variables in West Nile virus model. Available at: <https://github.com/desireeka93/WNV-Activity>.

Data S1. (separate file)

Code available at: https://github.com/desireeka93/WNV-Activity.
